# Increased preference for lysine over arginine in spike proteins of SARS-CoV-2 BA.2.86 variant and its daughter lineages

**DOI:** 10.1101/2024.11.11.622903

**Authors:** Anže Božič, Rudolf Podgornik

## Abstract

The COVID-19 pandemic offered an unprecedented glimpse into the evolution of its causative virus, SARS-CoV-2. It has been estimated that since its outbreak in late 2019, the virus has explored all possible alternatives in terms of missense mutations for all sites of its polypeptide chain. Spike protein of the virus exhibits the largest sequence variation in particular, with many individual mutations impacting target recognition, cellular entry, and endosomal escape of the virus. Moreover, recent studies unveiled a significant increase in the total charge on the spike protein during the evolution of the virus in the initial period of the pandemic. While this trend has recently come to a halt, we perform a sequence-based analysis of the spike protein of 2665 SARS-CoV-2 variants which shows that mutations in ionizable amino acids continue to occur with the newly emerging variants, with notable differences between lineages from different clades. What is more, we show that within mutations of amino acids which can acquire positive charge, the spike protein of SARS-CoV-2 exhibits a prominent preference for lysine residues over arginine residues. This lysine-to-arginine ratio increased at several points during spike protein evolution, most recently with BA.2.86 and its sublineages, including the recently dominant JN.1, KP.3, and XEC variants. The increased ratio is a consequence of mutations in different structural regions of the spike protein and is now among the highest among viral species in the *Coronaviridae* family. The impact of high lysine-to-arginine ratio in the spike proteins of BA.2.86 and its daughter lineages on viral fitness remains unclear; we discuss several potential mechanisms that could play a role and that can serve as a starting point for further studies.

## Introduction

Since its outbreak in late 2019, severe acute respiratory coronavirus 2 (SARS-CoV-2) has been undergoing numerous evolutionary changes which are reflected in changes in its rate of transmission, immune escape, and overall fitness [1]. The first significant change in the virus’ adaptation to humans has been observed about a year into the COVID-19 pandemic with the emergence of the first variant of concern (VOC) termed Alpha; soon thereafter, further VOCs such as Beta, Delta, and Omicron emerged and became dominant by outcompeting previous variants [1]. This process has led to the fairly recent emergence of Omicron BA.2.86 with enhanced antibody evasion [2] and the current prevalence of some of its descendant lineages, including JN.1, KP3, and XEC. Of the latter three, JN.1 shows a considerably higher infectivity and immune evasion at the expense of reduced binding to the angiotensin-converting enzyme 2 (ACE2) [3, 4]; KP.3, on the other hand, shows both strong immune evasion as well as increased receptor binding capability [5]. The XEC variant includes two new mutations compared to KP.3, which introduce potential glycosylation sites and thus further enhance immune evasion of the virus [6].

It has been estimated that, on average, any neutral single-nucleotide mutation in the SARS-CoV-2 genome has occurred ∼15000 independent times since its emergence, implying that the virus has already explored all possible alternatives in terms of missense mutations [7]. Variants emerged after late 2023 even appear to be incorporating reversions to residues found in other sarbecoviruses [8]. The mutational spectrum of SARS-CoV-2 is, however, highly uneven and shows preferences for certain mutations over others [9, 10], which can have important structural implications [11]. Spike protein of SARS-CoV-2—and its receptor-binding (RBD) and N-terminal (NTD) domains in particular—is the part of the virus that exhibits the greatest sequence variation [12]. Various specific mutations in the spike proteins of different SARS-CoV-2 variants have been linked to increased transmissibility of the virus and its binding to ACE2 and other receptors [13–19]. Some of these mutations create changes to and from different *ionizable amino acids* (amino acids whose residues carry either a positive or negative charge) and thus have the potential to impact electrostatic interaction of the spike protein with its environment [20–24]. Pawłowski [25, 26] observed a trend in the early stages of the pandemic in which mutations in ionizable amino acids acted to increase the *total charge* on the spike protein towards more positive values. This tendency, affecting the properties of the spike protein as a whole, has been confirmed by later studies on larger sets of SARS-CoV-2 lineages [27, 28], which have furthermore shown a plateauing of the total charge on the spike protein with the emergence of Omicron and subsequent variants [28–30]. Protein-wide changes in the total charge are especially relevant for non-specific electrostatic interactions of the spike protein with other charged macromolecules and macromolecular substrates in its environment [31–34]. For instance, the total charge on the RBD region of the spike protein has been shown to correlate with RBD–ACE2 affinity in the early VOCs of SARS-CoV-2 [35].

Even though the early changes in the total charge on the spike protein might have come to a halt [28–30], mutations that change the number and nature of ionizable amino acids continue to occur unabated. In the recently emerged lineage BA.2.86, for example, electrostatic changes have been shown to contribute to the immune evasion of the virus [36]. In this work, we perform a sequence-based analysis of the mutations in ionizable amino acids on the spike proteins of 2665 SARS-CoV-2 lineages that have emerged since the start of the pandemic. We find that the BA.2.86 variant and its daughter lineages show a notable increase in the preference for lysine residues over arginine residues, despite the fact that both amino acid types carry positive charge and should thus have a similar effect on the resulting electrostatic interactions. What is more, we observe that the ratio of lysine to arginine in the currently prevalent SARS-CoV-2 variants is among the highest seen in different viruses from the *Coronaviridae* family. We also demonstrate that the changes to this ratio have been consistently occurring despite the fact that mutations from lysine to arginine or vice versa typically do not have a significant effect on viral fitness, ACE2 binding affinity, or RBD expression. While the impact of the high lysine-to-arginine ratio in SARS-CoV-2 variants thus remains unclear, we provide several possible reasons that could explain it.

## Materials and methods

### Sequences, divergence, and date of emergence of SARS-CoV-2 lineages

Data collection of sequences from different SARS-CoV-2 lineages follows our approach described previously [28, 29]. In short, we use a list of SARS-CoV-2 Pango lineages from CoV-Lineages.org [37] (accessed 12. 12. 2024) to download SARS-CoV-2 genomic and protein data from NCBI Virus database [38] together with accompanying annotations. We define the date of emergence of a lineage as the earliest full record (i.e., year, month, and day) of isolate collection. Lineage divergence, defined through the number of mutations in the entire genome relative to the root of the phylogenetic tree (the start of the outbreak), is obtained from the global SARS-CoV-2 data available at Nextstrain.org [39] (accessed 16. 12. 2024), and only those entries with a genome coverage of *>* 99 % are selected. Finally, we retain the lineages whose downloaded fasta protein file is not empty, resulting in a total number of *N* = 2665 analyzed SARS-CoV-2 lineages. The final lists of analyzed lineages, the dates of their emergence, and their average divergence are available in OSF at https://osf.io/pfzbj/, reference number PFZBJ.

### Spike protein sequences of viruses from *Coronaviridae* family

As a point of comparison, we also examine the spike proteins of other viruses belonging to the *Coronaviridae* family. We focus on the *Orthocoronavirinae* subfamily, which is composed of four genera—*Alphacoronavirus, Betacoronavirus, Gammacoronavirus*, and *Deltacoronavirus*—and contains the majority of known coronaviruses [40]. Following the approach we use for SARS-CoV-2 sequences, we use the NCBI Virus database [38] (accessed 06. 01. 2025) to download the spike protein sequences and annotations for different viruses belonging to *Orthocoronavirinae*. We retain only the entries with complete sequences without any ambiguous characters. The final dataset contains 64 viruses from genus *Alphacoronavirus*, 36 viruses from genus *Betacoronavirus* (excluding SARS-CoV-2), 8 viruses from genus *Gammacoronavirus*, and 17 viruses from genus *Deltacoronavirus*. The list of analyzed coronaviruses, their spike protein sequences, and annotations are available in OSF at https://osf.io/pfzbj/, reference number PFZBJ.

### Ionizable amino acids

Typically, six different amino acids can acquire charge [41, 42]; aspartic (ASP) and glutamic (GLU) acid as well as tyrosine (TYR) acquire negative charge while arginine (ARG), lysine (LYS), and histidine (HIS) acquire positive charge. We neglect cysteine (CYS) as it is usually not considered to be an acid [41, 42]. We use Biopython [43] to parse the dowloaded protein fasta files and count the number of ionizable amino acids on the spike proteins of different SARS-CoV-2 lineages. The results of this analysis are available in OSF at https://osf.io/pfzbj/, reference number PFZBJ.

To compare the relative change in the number of ionizable amino acids on the spike proteins of different lineages compared to the wild-type (WT) version (lineage B), we define

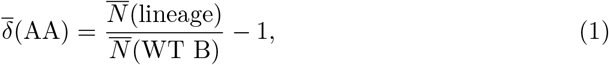

where 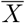 denotes the average over all spike protein sequences included under each lineage. This measure allows for an easier comparison of the patterns of change in the number of ionizable amino acids in a large number different SARS-CoV-2 lineages.

### Effects of mutations on SARS-CoV-2 fitness

To study the effect of mutations of ionizable amino acids on different aspects of SARS-CoV-2 fitness, we use publicly available datasets from several studies. Foremost, we use the data from Ref. [10], where the effect of mutations on viral fitness was estimated through the logarithm of observed vs. expected mutations in a large dataset of SARS-CoV-2 sequences (https://github.com/jbloomlab/SARS2-mut-fitness; accessed 24. 12. 2024). We furthermore use the data from Ref. [18] which contains the effects of all single amino-acid mutations on the binding of the spike protein to ACE2 as well as on RBD expression—a proxy for folding and stability—in the context of several SARS-CoV-2 variants (https://github.com/jbloomlab/SARS-CoV-2-RBD_DMS_variants/; accessed 24. 12. 2024).

## Results

### Lineage-specific changes in the number of ionizable amino acids on the spike protein

Of the six ionizable amino acid types that typically carry charge, three of them are negatively charged (TYR, ASP, and GLU), and three of them are positively charged (ARG, LYS, and HIS). Missense mutations involving these amino acids can thus either *(i)* convert a neutral amino acid to a charged one or vice versa; *(ii)* convert a positively charged amino acid to a negatively charged one (o vice versa); or *(iii)* convert between different amino acid types that otherwise carry the same charge. Figure 1 shows the average relative change in the number of ionizable amino acids on the spike proteins of different SARS-CoV-2 lineages compared to WT lineage B. First thing we observe is that these mutations lead to distinct profiles of changes as lineage divergence increases, and we have already shown previously that the clustering of lineages based on these patterns of change broadly follows their phylogenetic division into clades [29]. The currently dominant set of lineages, which includes JN.1, KP.3, and XEC and traces their origin to the BA.2.86 variant, presents a very distinct cluster characterized by a notable increase in the number of LYS residues and a decrease in the number of ARG and GLU residues (highlighted box in Fig 1). The pattern of change in this cluster is very distinct from previous variants, including, for instance, XBB and other recombinant variants, which show a high relative enrichment and variability in the number of HIS residues but less drastic changes to the number of ARG and LYS residues. The recently emerged variants also exhibit large changes compared to the early VOCs such as Beta and Delta, where the increase towards overall positive charge on the spike protein has first been observed [25, 26].

**Fig 1.**
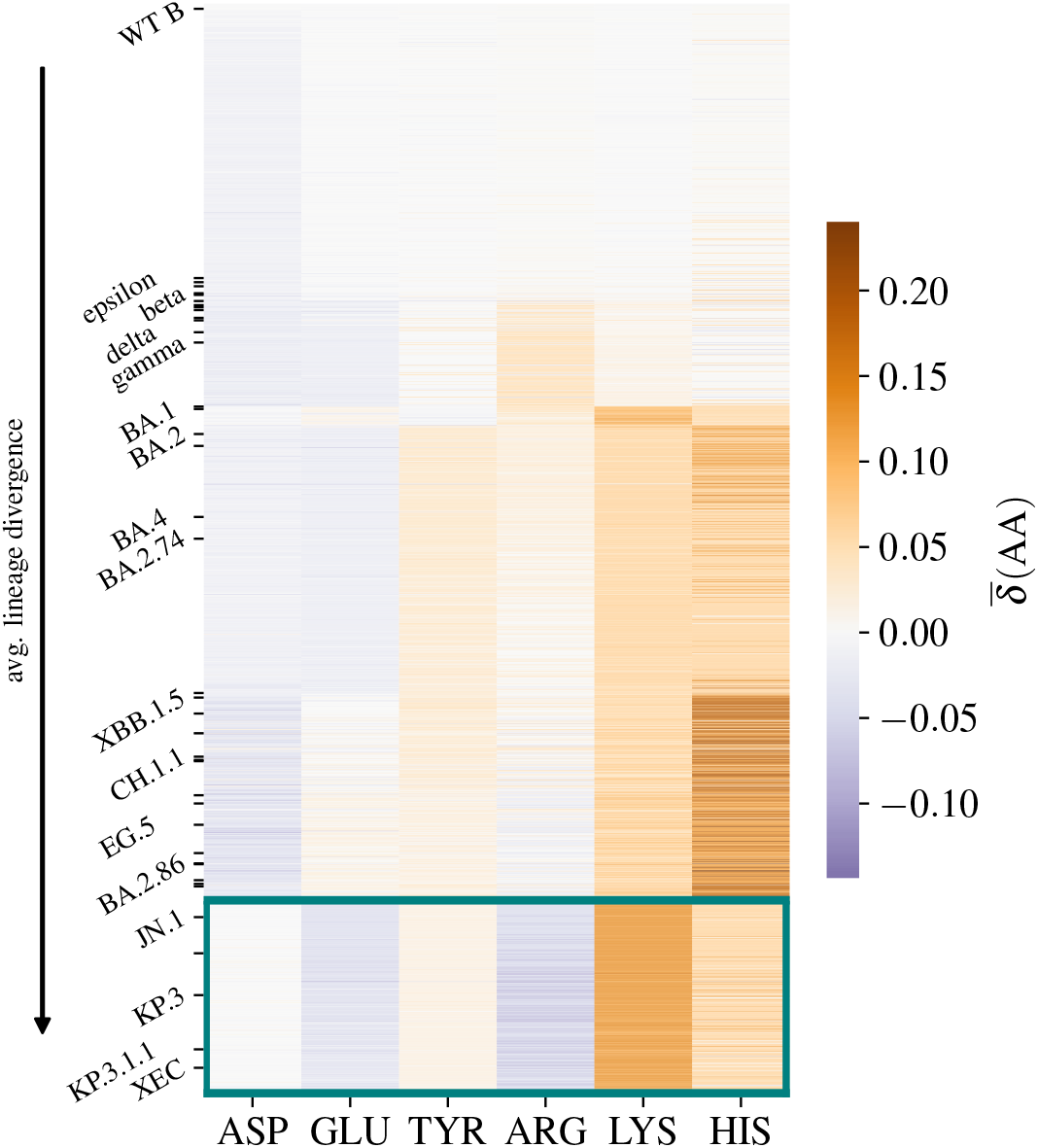
Average change in the number of ionizable amino acids on the spike protein of SARS-CoV-2 lineages. Shown is the average change 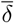 (AA) [Eq (1)] in the number of negatively charged (ASP, GLU, and TYR) and positively charged (ARG, LYS, and HIS) amino acids on the spike protein of 2665 different SARS-CoV-2 lineages compared to WT lineage B. A relative change of 10% typically corresponds to an addition or deletion of ∼5 residues, depending on the amino acid in question (see also Fig 2). Lineages are ordered according to their (average) divergence from WT B. Highlighted box includes the currently dominant set of lineages, including JN.1, KP.3, KP.3.1.1, and XEC, all showing similar patterns of change in the number of ionizable amino acids.

We can also look into the relative changes in the number of ionizable amino acids on three of the main structural domains of the spike protein, the N-terminal domain (NTD), RBD, and S2 domain. While the changes in the number of ionizable amino acids act globally to influence the total charge carried by the spike protein, their distribution within the structural domains is far from uniform, and each domain can be characterized by a particular pattern of changes (Figure in S1 Fig). The spike protein RBD is known to be positively charged for a broad range of pH values [44, 45], and the evolutionary changes in ionizable amino acids in this region have generally acted to increase the amount of positive charge, in particular by increases in both the number of LYS and HIS residues. The NTD, on the other hand, shows a decrease in the number of ARG and HIS residues which act to decrease the amount of positive charge; this corresponds with previous studies showing that this region underwent several changes during SARS-CoV-2 evolution which made it net negative [46]. Lastly, the stalk part of the spike protein—the S2 domain—is evolutionary very conserved and is known to be overall negatively charged [44, 45]; there, we observe a pattern of change virtually conserved since the emergence of Omicron variant, at which point the number of LYS, HIS, and TYR residues have increased.

### Lysine-to-arginine ratio has increased several times during spike protein evolution

Changes in the number of ionizable amino acids on the spike protein thus continue to occur unabated despite the fact that the early increase in the total charge on the spike protein [25–28] had essentially stopped with the emergence of Omicron and later lineages [28, 29]. Even if they do not impact the total charge, mutations of amino acids which carry charge can still have significant influence on both the properties of the spike protein as well as on the overall viral fitness [17, 47]. A particularly notable trend in the recently emerged SARS-CoV-2 lineages is the relative decrease in the number of ARG residues (Fig 2a) and a simultaneous increase in the number of LYS residues (Fig 2e).

**Fig 2.**
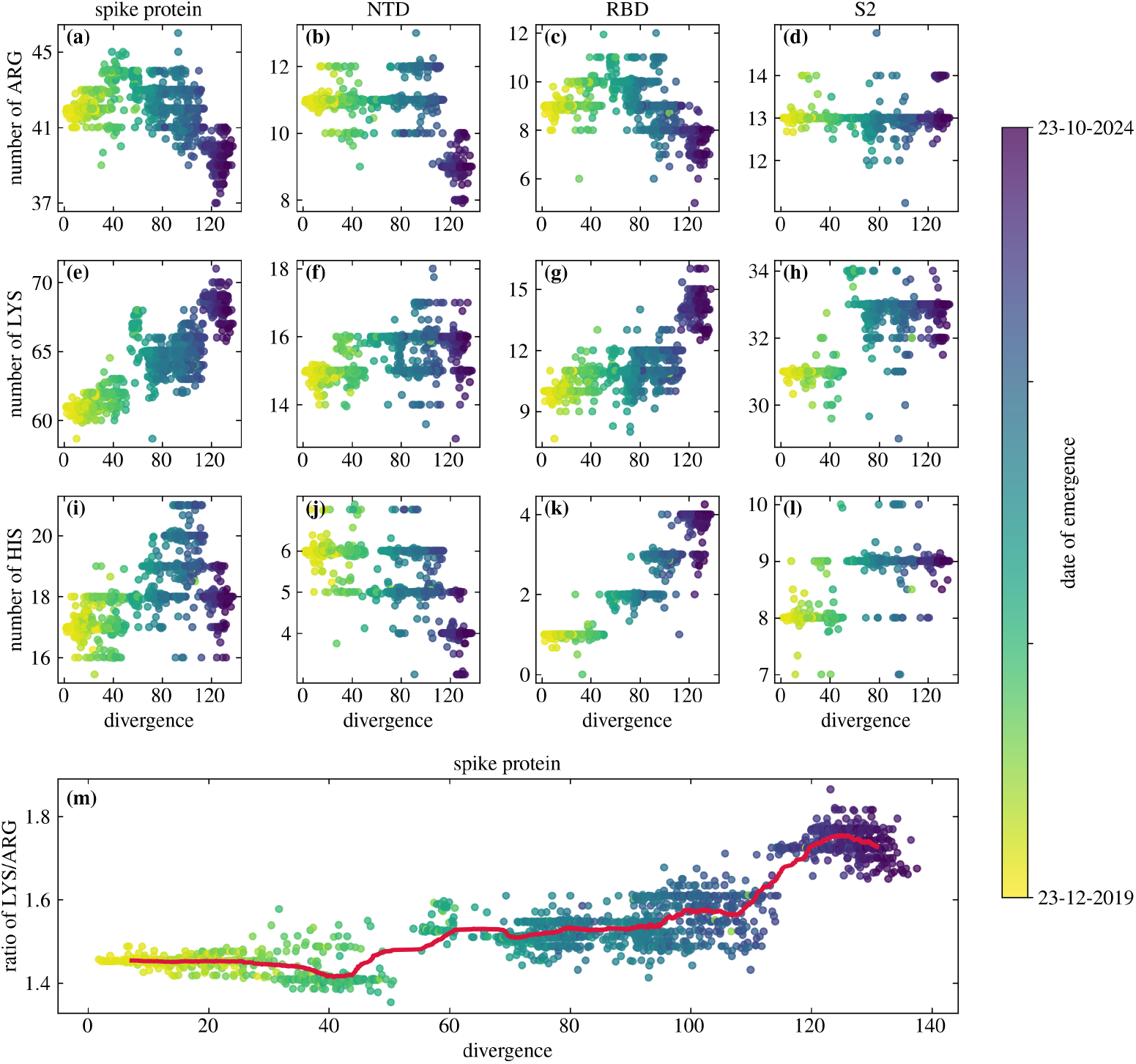
Evolutionary changes in the number of arginine, lysine, and histidine residues on the spike protein of SARS-CoV-2 and its different regions. Shown is the change in the number of **(a)–(d)** ARG, **(e)—(h)** LYS, and **(i)–(l)** HIS residues, both on the entire spike protein (first column) as well as on the N-terminal domain (NTD; second column), receptor-binding domain (RBD; third column) and S2 region (fourth column) alone. Panel **(m)** shows the ratio of LYS to ARG on the entire spike protein as a function of the (average) divergence. Each point in the panels represents one of the 2665 different SARS-CoV-2 lineages analyzed. Thick line in panel (m) shows a rolling average of the LYS/ARG ratio with lineage divergence (window size contains 100 lineages).

The increase in the number of LYS residues—from ∼60 at the start of the COVID-19 pandemic to ∼70 with the most recently emerged lineages—stands out in particular and is the main reason for the observed increase in the LYS/ARG ratio (Fig 2m). This ratio has increased twice during the timespan of the pandemic, first with the early Omicron lineages—where also the total charge on the spike protein reached its peak [28, 29]—and next with the recent emergence of BA.2.86 and its daughter lineages. In contrast to LYS and ARG, HIS, another positively charged amino acid, shows larger variability with lineage divergence and no clear evolutionary trend (Fig 2i).

Looking further into the variation in the number of LYS and ARG residues in specific structural domains of the spike protein shows that these changes have not occurred simultaneously throughout the protein (Figs. 2 and 3 and Figure in S2 Fig).

**Fig 3.**
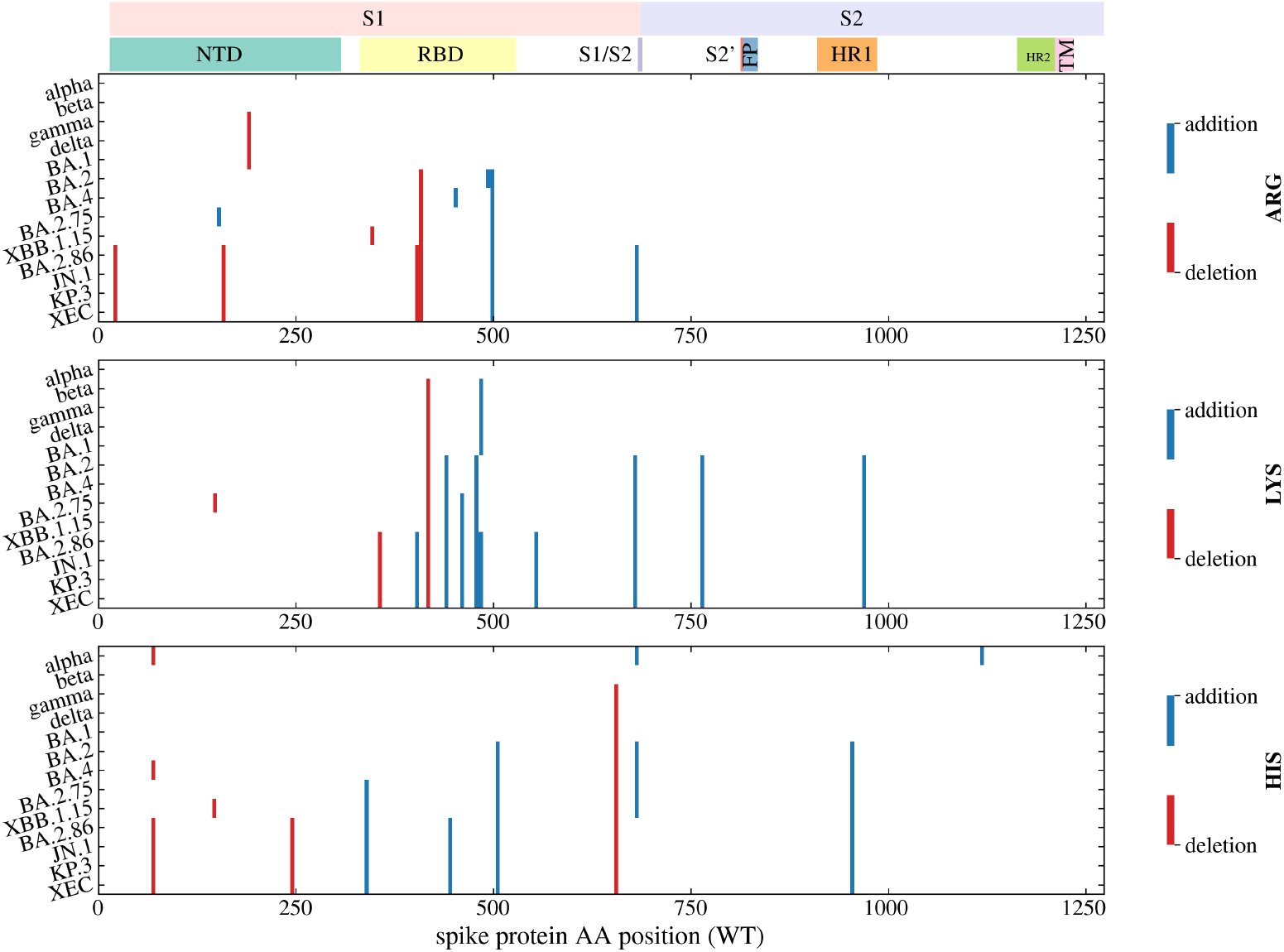
Characteristic mutations in arginine, lysine, and histidine residues in the spike proteins of select SARS-CoV-2 variants. Shown are those substitutions to (“addition”) or from (“deletion”) three positively-charged amino acid types (ARG, LYS, and HIS) that occur in at least 70% of the spike protein sequences of a given variant (data from CoV-Spectrum.org [48]; accessed 08. 01. 2025).

The first notable increase in the LYS/ARG ratio has mainly occurred in the S2 domain (Figure in S2 Fig) and is largely due to the addition of several new LYS residues (Figs 2h and 3). The second notable increase occurred due to further new substitutions to LYS residues and from ARG residues in the RBD region of the spike protein (Figs 2c and 2g and Figure in S2 Fig). This is further illustrated in Fig 3, which shows the characteristic substitutions in the three positively charged amino acid types of select SARS-CoV-2 variants. Substitutions to and from LYS and ARG residues most often occur in the RBD region of the spike protein, and several of these mutations are characteristic of only the BA.2.86 variant and its descendant lineages. The NTD region, on the other hand, exhibits only a few substitutions from ARG, which is in line with observations that this region tends to acquire a more negative charge [46].

### Betacoronaviruses have the highest lysine-to-arginine ratio in *Coronaviridae*

To better understand the importance of the observed increase in the LYS/ARG ratio in emerging SARS-CoV-2 variants, we also take a look at this ratio in the spike proteins of other viral species from the *Coronaviridae* family. As Fig 4 shows, viruses in the *Betacoronavirus* genus have a significantly higher ratio of LYS/ARG compared to viruses from *Alphacoronavirus* and *Deltacoronavirus* genera (Brunner-Munzel test, *p <* 0.0001). Even more interesting is the fact that the mean value of LYS/ARG ratio in betacoronaviruses (LYS*/*ARG ≈ 1.34) corresponds well with the minimum value of this ratio obtained by any SARS-CoV-2 lineage (LYS*/*ARG ≈ 1.35; lower dashed line in Fig 4). Moreover, while a number of betacoronaviruses have the value of this ratio above the mean—with the maximum in our dataset achieved by the spike protein of Hedgehog coronavirus 1 (LYS*/*ARG ≈ 1.68)—this is still far from the maximum value achieved by any SARS-CoV-2 lineage thus far (LYS*/*ARG ≈ 1.86; upper dashed line in Fig 4). These observations show that upon emergence, the initial LYS/ARG ratio of the SARS-CoV-2 spike protein was around the average value characteristic of betacoronaviruses; however, this ratio has increased during its evolution to values not found in any of the viruses in the dataset of *Coronaviridae* we examined.

**Fig 4.**
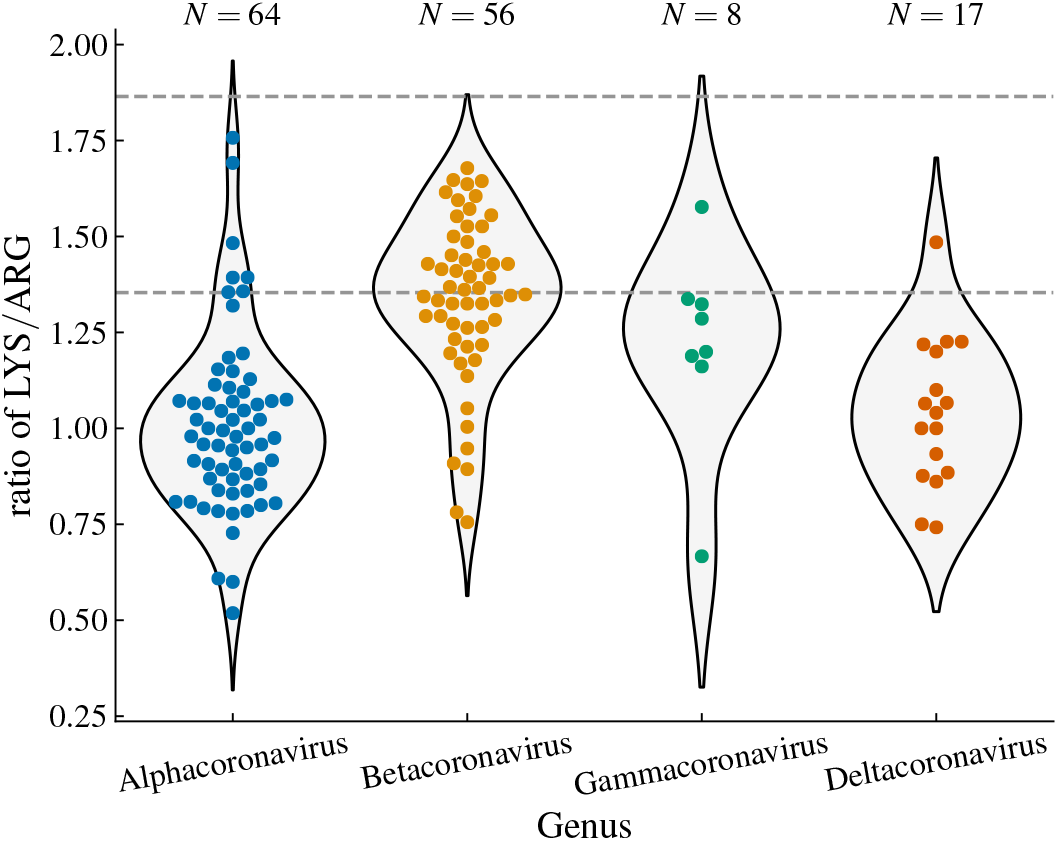
Ratio of lysine to arginine in the spike proteins of different viral species belonging to *Coronaviridae* family. Shown are the violin plots as well as individual data points for viruses belonging to the four genera in the *Orthocoronavirinae* subfamily. Each data point represents a species with a unique name in the NCBI Virus database [38], with the ratio averaged over all database entries for that species. Dashed lines show the minimum and maximum value of the ratio that are obtained by the analyzed SARS-CoV-2 lineages (see Fig 2).

### Replacing spike protein lysines with arginines does not significantly impact viral fitness

The increase in the LYS/ARG ratio during the evolution of SARS-CoV-2 is particularly interesting since it does not in any obvious way impact the overall positive charge on the spike protein, and we need to explore other potential reasons for this change. We can draw on previous studies which have explored how mutations impact different aspects of SARS-CoV-2 fitness [10, 49] to examine whether ARG↔LYS mutations in the spike protein come with some consequence for viral fitness. If so, this would then imply that the mutations of an amino acid to LYS comes with a benefit which would not be present if the amino acid were mutated to ARG instead.

Change in viral fitness was for instance estimated by Bloom and Neher [10] as the logarithm of actual vs. observed mutation counts in publicly available SARS-CoV-2 sequences. Fig 5 shows this change in viral fitness if ARG residues are replaced by LYS residues and vice versa. In the spike protein (Fig 5a), the change of LYS to ARG typically does not come with a change in viral fitness. Changing ARG to LYS, on the other hand, sometimes leads to a larger decrease in viral fitness, implying that certain ARG residues are crucial to it. This difference is, interestingly, not bound to the spike protein but can be observed even more strongly when we consider these mutations in all other protein-coding genes (Fig 5b). On the other hand, if one considers mutations from ARG to any other residue or from any residue to LYS, this effect disappears (Figure in S3 Fig), in line with a previous report that most (but not all) mutations have similar effects on the spike proteins of different SARS-CoV-2 variants [17].

**Fig 5.**
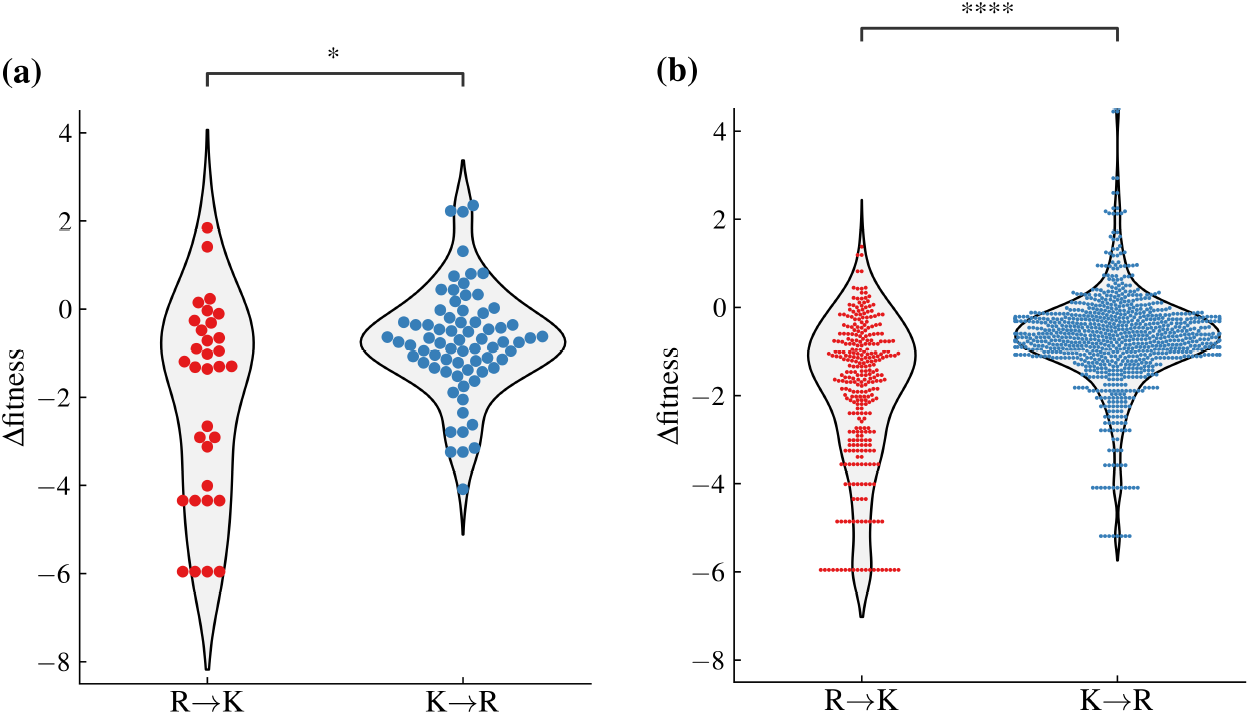
Influence of mutations between arginine and lysine on SARS-CoV-2 fitness. Viral fitness is estimated as the logarithm of actual vs. observed mutation counts in all publicly available SARS-CoV-2 sequences as of March 2023; data are taken from Ref. [10]. Shown are the mutations in **(a)** the spike protein and in **(b)** all other protein-coding regions. Mutations from arginine to lysine and from lysine to arginine have different effect on viral fitness in both datasets (Brunner-Munzel test; (a) *p* = 0.018 and (b) *p <* 0.0001).

A similar difference between the effects of LYS →ARG and ARG→ LYS substitutions in the spike protein can be observed when we utilize the data of Taylor *et al*. [49], who measured the change in the binding affinity of the spike protein to ACE2 or in the expression of RBD upon mutations in select SARS-CoV-2 variants (Figure in S4 Fig). While changing any LYS residue to an ARG residue has little effect on any of the two measures, this is conversely not always the case when an ARG residue is changed to a LYS residue. That the effects of LYS↔ARG mutations should be similar between the two datasets is nor surprising, as another study [17] already showed that the effects of mutations on cell entry were fairly well correlated with the effects of amino-acid mutations on viral fitness. Since these data imply LYS→ARG mutations do not influence the obvious markers of SARS-CoV-2 fitness, there should thus be yet another reason still why mutations to LYS nonetheless seem to be preferred over mutations to ARG.

## Discussion

Studies have shown that viral proteins generally tend to contain more ARG than LYS [50] and that an infected cell absorbs far more ARG from the culture medium than an uninfected one [51]. This led to the idea of diet restriction of ARG-rich foods during viral illnesses [52] and supplementation with LYS, its antagonist, which attenuates the growth-promoting effect of ARG [53]. There is a competitive antagonism between LYS and ARG, and if a cell gets saturated in one of the two, its absorption slows and leaves the other free to be more absorbed [54]. Some studies have shown positive results of ARG depletion on treating viral infections [52, 54], including SARS-CoV-2 [55]; however, a recent study by Rees *et al*. [56] suggests this might potentially exacerbate the effects of the infection by SARS-CoV-2 instead. Moreover, a study by Melano *et al*. [57] has demonstrated that it is in fact LYS restriction that can attenuate SARS-CoV-2 infection, which appears to be in line with our observation that SARS-CoV-2 spike proteins of recent lineages increasingly prefer LYS over ARG (Fig 2). This is also a common feature we observe in other coronavirus spike proteins as well (Fig 4) and which stands in contrast to what is typically observed in other viruses [50].

The increasing ratio of LYS/ARG in spike proteins of emerging SARS-CoV-2 variants, high even in comparison to other betacoronaviruses, cannot be simply related to codon usage bias. While it is true that ARG is coded for by six codons and LYS by only two, codon usage bias in SARS-CoV-2 and other coronaviruses suggests that CpG codons are suppressed, with only two out of the six ARG codons being predominantly in use [58, 59]. However, the reasons behind the preference for LYS over ARG might be related to their physicochemical properties. For instance, in the membrane, LYS residues readily deprotonate while ARG residues maintain their charge [60, 61] and can more easily disrupt and permeabilize lipid membranes. ARG also forms a larger number of electrostatic interactions with the surrounding compared to LYS [62, 63]. Thus, preference for LYS over ARG could favourably influence the electrostatic interactions involving SARS-CoV-2 spike protein in specific environments [31, 64, 65].

Studies have also shown that preference for LYS over ARG can influence protein structural stability and folding [62, 66, 67], and a computational structural analysis of SARS-CoV-2 spike protein revealed that three specific mutations of asparagine (N) to lysine in the central core region (N764K, N856K, and N969K) contribute to a preference for the alteration of the spike protein conformations [68, 69]. Furthermore, the ratio of ARG to LYS is a known factor in determining protein solubility and aggregation, with LYS being enriched relative to ARG in many of the more soluble proteins [67, 70–73]. However, a quick comparison of the spike protein solubilities in SARS-CoV-2 variants, predicted from their sequence using `protein-sol` [74], shows no difference between them despite their different LYS/ARG ratios. Even more, such an analysis performed on spike proteins of different coronaviruses indicates that spike protein solubility in general does not change much once the LYS/ARG ratio is sufficiently large (LYS*/*ARG ≳ 1.2; Figure in S5 Fig). While a more detailed analysis performed on spike protein structures would be needed to obtain more reliable estimates [75], these results indicate that the observed increase in LYS/ARG ratio in spike proteins of SARS-CoV-2 variants is unlikely to modify their solubility.

An important difference between ARG and LYS is that the latter is far more prone to post-translational modifications (PTMs) [76, 77]. While both amino acids can undergo methylation, LYS residues can also undergo ubiquitination, which can regulate the association of the spike protein with ACE2 [78], and acetylation, which can contribute to a range of virus–virus and virus–host interactions by, e.g., regulating interactions between proteins and membranes and enabling generation of novel protein-binding recognition surfaces [79–83]. What is more, LYS acetylation is a reversible process that leads to neutralization of the position’s positive electrostatic charge [84] and can contribute to structural changes [85, 86]. While PTMs have been mostly studied on non-spike proteins of SARS-CoV-2 [87], a computational study by Liang *al*. [85] identified 87 PTM sites from 5 major modifications on the spike protein of SARS-CoV-2 variant Alpha, of which the largest number of modified sites corresponded to glycosylation (39 residues) followed by acetylation (21 residues). The influence of these PTMs was studied via “mutagenesis” amino acid substitution rules, which in the case of LYS and ARG residues corresponds to the substitution LYS↔ARG. This study [85] nonetheless did not observed any marked difference between the spike protein’s unmodified and modified structures, especially for the functional regions in the S1 and S2 domains. Since Alpha variant is situated at the very beginning of SARS-CoV-2 evolution, more insight is needed into PTMs in the spike proteins of recently emerged variants whose profiles of ionizable amino acids are, as we have seen, characterized by numerous new LYS residues.

Lastly, it is possible that the observed increase in the LYS/ARG ratio on the spike proteins of emerging SARS-CoV-2 variants—which does not increase the overall charge on the spike protein nor is easily related to any obvious markers of viral fitness—can be related to the attenuation of SARS-CoV-2 as a part of its adaptation to the new host. Upon cross-species transmission of a virus, its virulence often becomes markedly more lethal [88], which is also the case with SARS-CoV-2 [89]. In the initial evolution of SARS-CoV-2, transmissibility and virulence could increase in parallel. This trajectory changed with the emergence of Omicron whose increased transmission (due to, in part, changes in ionizable amino acids [90]) came at the expense of a decrease in virulence [89]. This is characteristic of the virulence-transmission trade-off hypothesis [88, 91], which predicts the selection of phenotypes with intermediate viral fitness. Such a relationship is, however, difficult to establish [88], especially in the light of the complex interplay of SARS-CoV-2 biology in the context of changing immunity due to both vaccination, antiviral drugs, and prior infection [1, 89].

## Conclusion

By examining the changes in the number of ionizable amino acids in the spike protein sequences of 2665 SARS-CoV-2 variants that have emerged since the start of the COVID-19 pandemic, our study identified an increasing ratio of LYS to ARG residues in the spike protein. This is a consequence of mutations that have been occurring since the early Omicron variants and which have seen a recent uptick with BA.2.86 and its daughter lineages. We have shown that these changes have been occurring throughout the spike protein, albeit not in all structural domains simultaneously. Comparing the observed LYS/ARG ratios with those of other coronaviruses, we observed that even though this ratio is in general higher in betacoronaviruses compared to other genera, the values it reached in the recently emerged SARS-CoV-2 variants are not seen in any other species in *Orthocoronavirinae*. Combined with observations from previous studies [10, 49], we also showed that the choice of LYS over ARG in the mutations found in emerging lineages most often does not come with a significant benefit associated with viral fitness, binding with the ACE2 receptor, or the expression of RBD. While the reasons behind the increase in the LYS/ARG ratio thus remain an open question, we have outlined several potential mechanisms that could play a role and that can hopefully serve as a starting point for further studies.

## Acknowledgments

A.B. acknowledges support from Slovenian Research Agency (ARIS) under contracts no. P1-0055 and no. J1-60002. R.P. acknowledges support from National Natural Science Foundation of China (NSFC) [Key Project 12034019]. There was no additional external funding received for this study.

## Supporting Information

**Fig S1.**
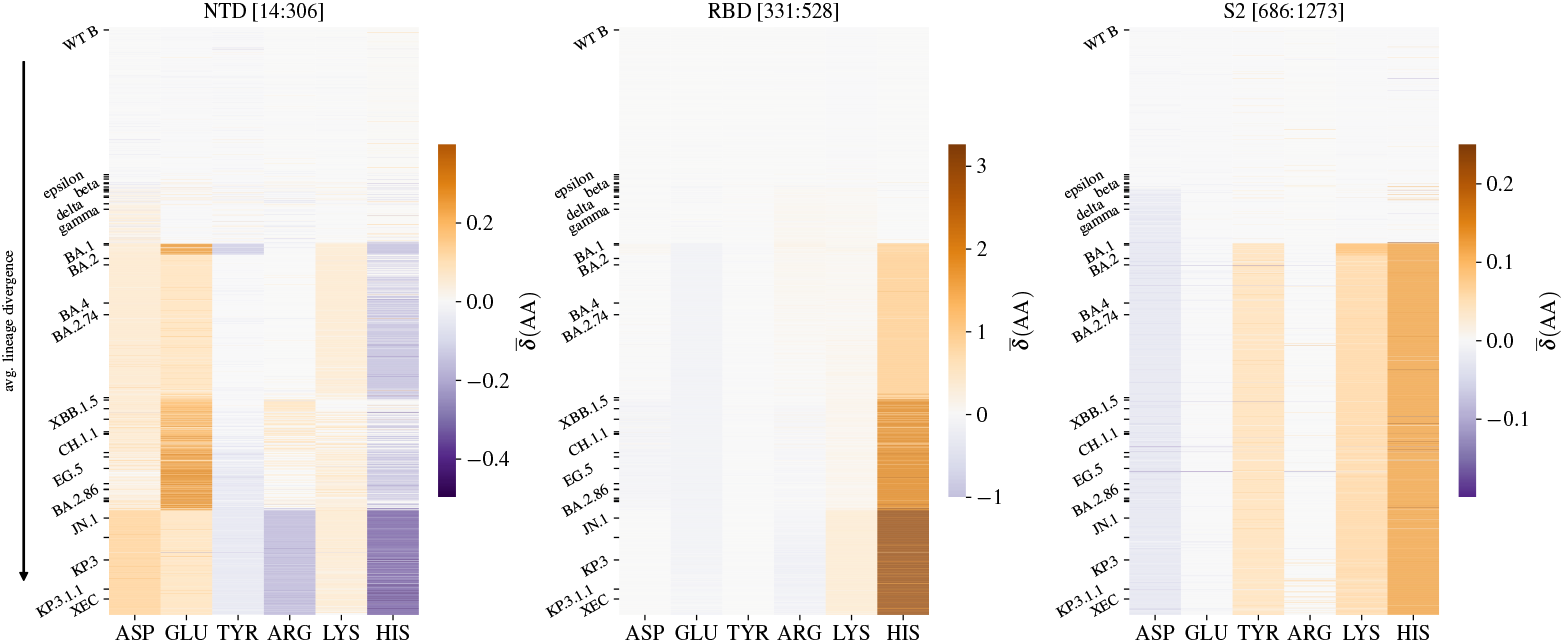
Average change in the number of ionizable amino acids on different structural regions of the spike protein of SARS-CoV-2 lineages. Shown are the changes in the N-terminal domain (NTD), receptor-binding domain (RBD), and S2 domain of the spike protein. Region boundaries are taken from Ref. [92] and are determined with respect to the WT SARS-CoV-2 spike protein sequence. Panels show the average change 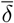(AA) [Eq (1)] in the number of negatively charged (ASP, GLU, and TYR) and positively charged (ARG, LYS, and HIS) amino acids of 2665 different SARS-CoV-2 lineages compared to WT lineage B. Lineages are ordered according to their (average) divergence from WT B.

**Fig S2.**
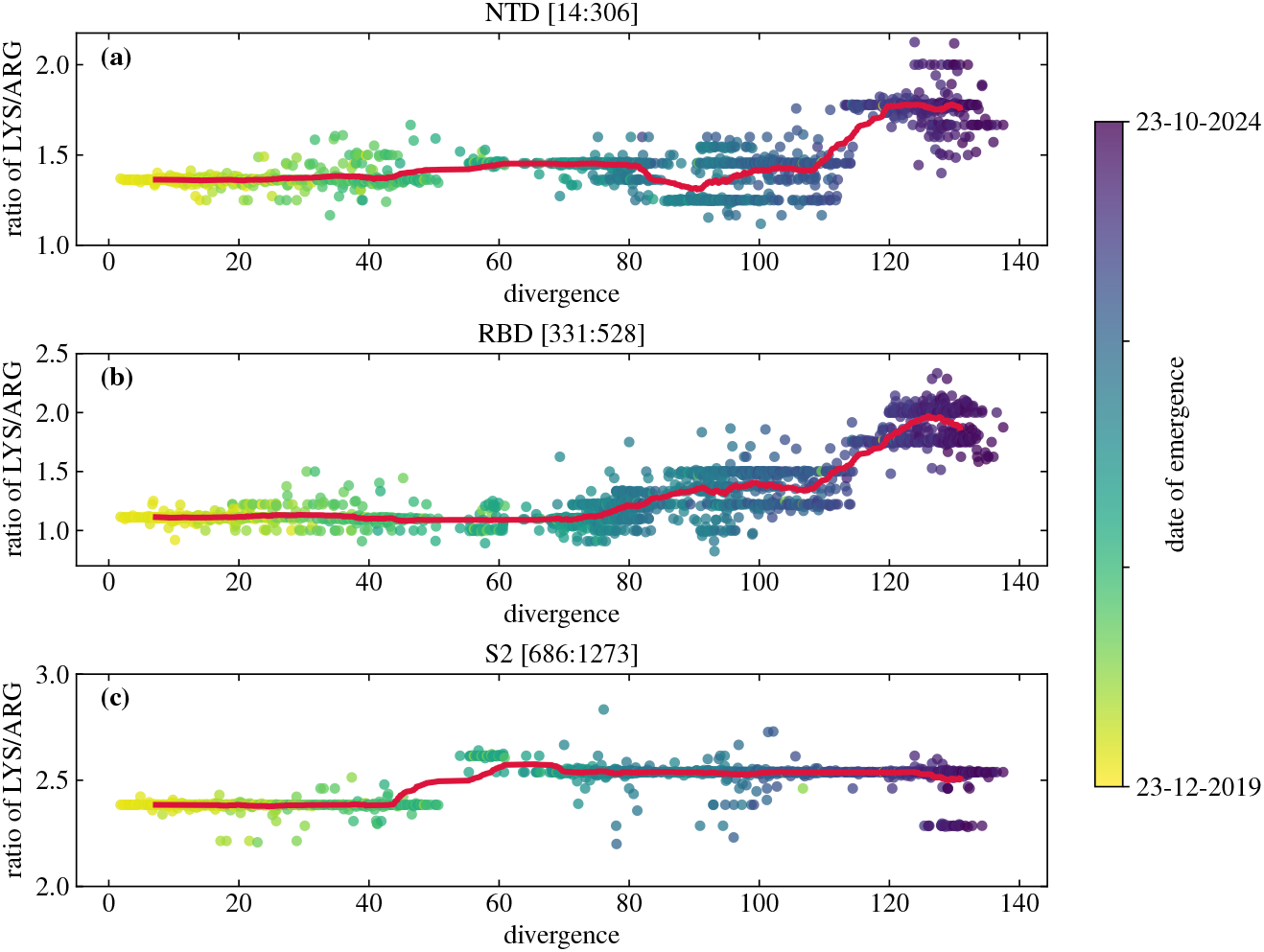
Evolutionary changes in the ratio of lysine to arginine on different structural regions of the spike protein of SARS-CoV-2 lineages. Shown are the changes in the LYS/ARG ratio in the **(a)** NTD, **(b)** RBD, and **(c)** S2 domain as a function of the (average) lineage divergence. Each point in the panels represents one of the 2665 different SARS-CoV-2 lineages analyzed. Thick lines show a rolling average of the LYS/ARG ratio with lineage divergence (window size contains 100 lineages).

**Fig S3.**
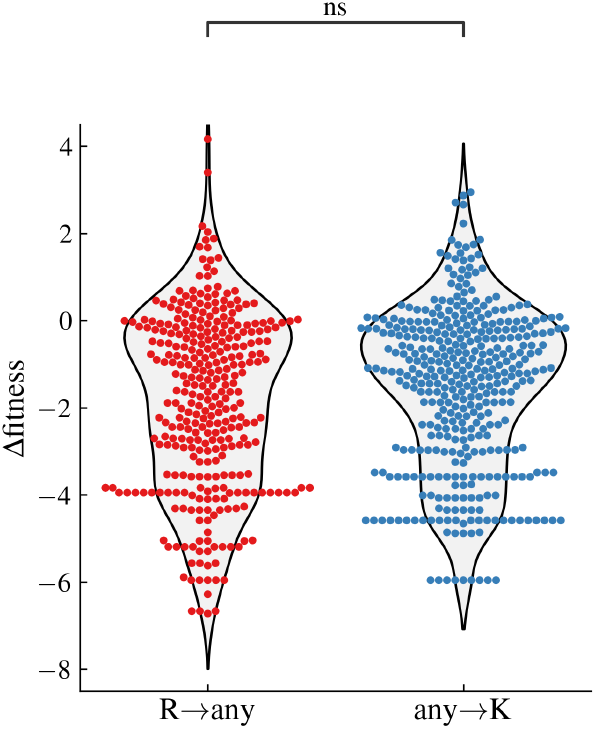
Influence of arginine and lysine mutations on SARS-CoV-2 fitness. Viral fitness is estimated as the logarithm of actual vs. observed mutation counts in publicly available SARS-CoV-2 sequences as of March 2023; data are taken from Ref. [10]. Compared are the mutations in the spike protein from arginine to any amino acid (R → any) and from any amino acid to lysine (any → K). These two types of mutations do not differ significantly in their effect on viral fitness (Brunner-Munzel test, *p* = 0.06).

**Fig S4.**
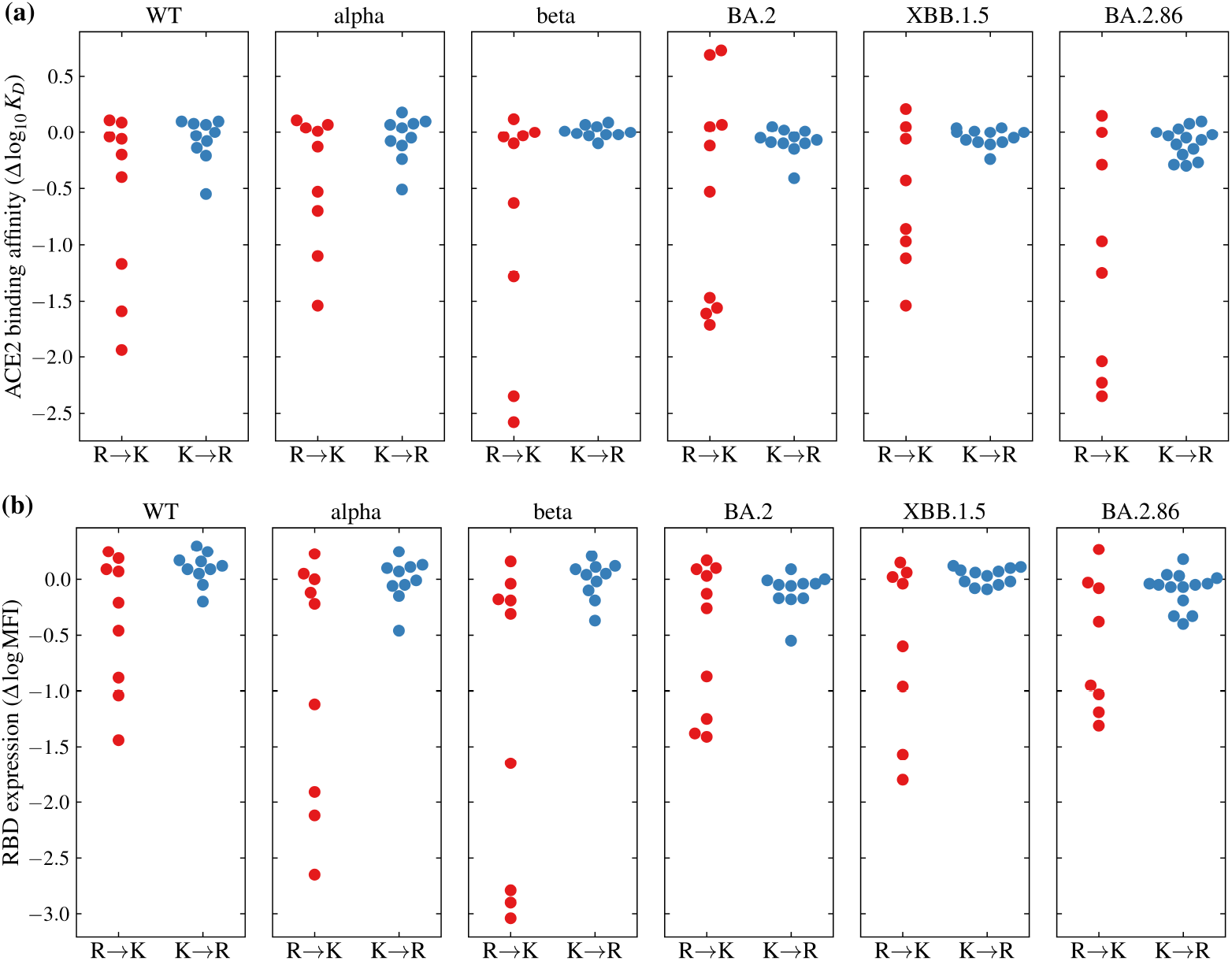
Influence of mutations between arginine and lysine on different aspects of SARS-CoV-2 fitness. Shown are the effects of spike protein mutations from ARG to LYS and vice versa on **(a)** ACE2 binding affinity and **(b)** RBD expression of select SARS-CoV-2 variants. Data are taken from Ref. [49].

**Fig S5.**
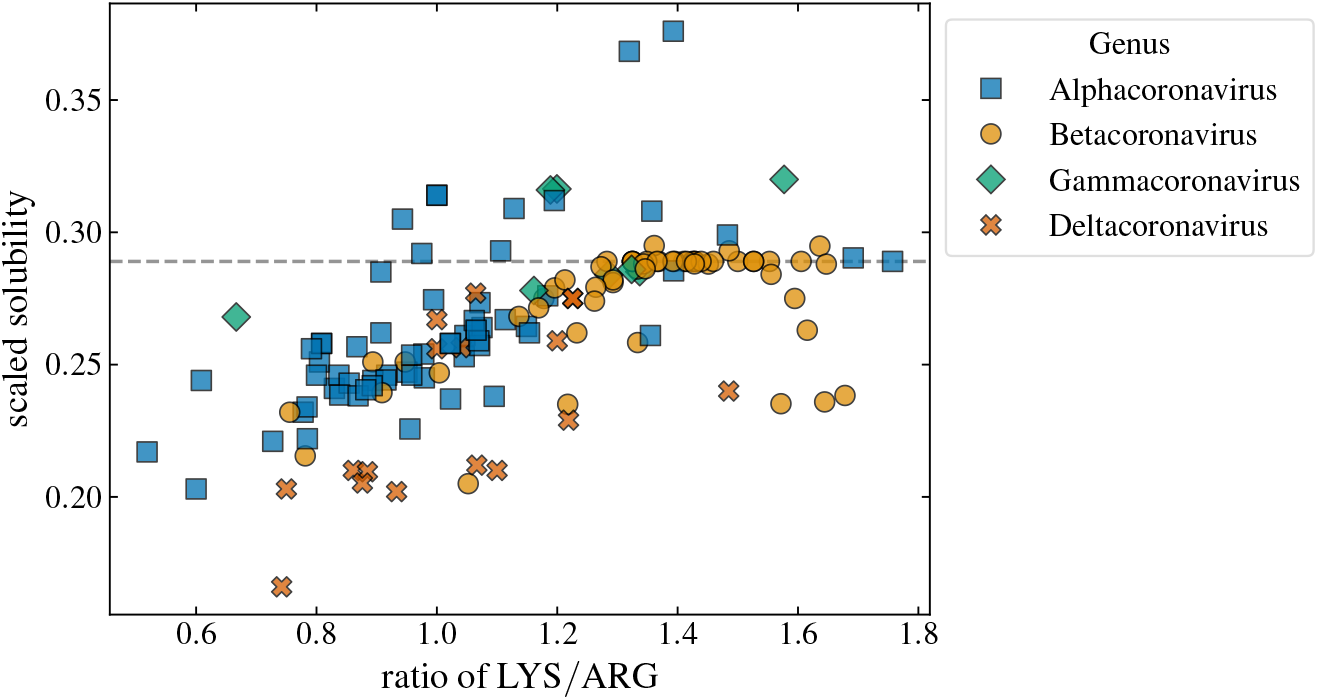
Ratio of lysine to arginine in spike proteins of different coronaviruses compared to their predicted (scaled) solubility. Dataset of viruses from the four genera of *Orthocoronavirinae* subfamily is the same as in Fig 4 in the main text. Scaled solubility was predicted from spike protein sequences using `protein-sol` [74]. Dashed line shows the predicted scaled solubility of 0.289, which is predicted to be the same for the entire dataset of SARS-CoV-2 spike protein sequences.

